# Upper respiratory microbial communities of healthy populations are shaped by niche and age

**DOI:** 10.1101/2024.04.14.589416

**Authors:** Susan Zelasko, Mary Hannah Swaney, Shelby Sandstrom, Timothy C. Davenport, Christine M. Seroogy, James E. Gern, Lindsay R. Kalan, Cameron R. Currie

**Affiliations:** Department of Bacteriology, University of Wisconsin-Madison, Madison, Wisconsin, USA; Microbiology Doctoral Training Program, University of Wisconsin-Madison, Madison, Wisconsin, USA; Department of Medical Microbiology and Immunology, School of Medicine and Public Health, University of Wisconsin-Madison, Madison, Wisconsin, USA; Department of Pediatrics, University of Wisconsin School of Medicine and Public Health, Madison, WI, USA; Department of Medicine, University of Wisconsin School of Medicine and Public Health, Madison, WI, USA; M.G. DeGroote Institute for Infectious Disease Research, David Braley Centre for Antibiotic Discovery, Department of Biochemistry and Biomedical Sciences, McMaster University, Hamilton, Ontario, Canada

## Abstract

**Background:** Alterations in upper respiratory microbiomes have been implicated in shaping host health trajectories, including by limiting mucosal pathogen colonization. However, limited comparative studies of respiratory microbiome development and functioning across age groups have been performed. Herein, we perform shotgun metagenomic sequencing paired with pathogen inhibition assays to elucidate differences in nasal and oral microbiome composition and functioning across healthy 24-month-old infant (n=229) and adult (n=100) populations.

**Results:** We find that beta diversity of nasal and oral microbiomes varies with age, with nasal microbiomes showing greater population-level variation compared to oral microbiomes. Infant microbiome alpha diversity was significantly lower across nasal samples and higher in oral samples, relative to adults. Accordingly, we demonstrate significant differences in genus- and species-level composition of microbiomes between sites and age groups. Antimicrobial resistome patterns likewise varied across body sites, with oral microbiomes showing higher resistance gene abundance compared to nasal microbiomes. Biosynthetic gene clusters encoding specialized metabolite production were found in higher abundance across infant oral microbiomes, relative to adults. Investigation of pathogen inhibition revealed greater inhibition of gram-negative and gram-positive bacteria by oral commensals, while nasal isolates had higher antifungal activity.

**Conclusions:** In summary, we identify significant differences in the microbial communities inhabiting nasal and oral cavities of healthy infants relative to adults. These findings inform our understanding of the interactions impacting respiratory microbiome composition and functioning, with important implications for host health across the lifespan.

## Introduction

The nasal and oral cavities are the first points of contact with the external environment and serve as gateways to the lower respiratory and gastrointestinal tracts. Inhaled air containing microbes passes through the nares before entering the nasal cavity that connects to the oropharynx and distal pulmonary sites via the nasopharynx [1]. Microbes entering the oral cavity pass to the digestive tract by swallowing and to the airways through micro-aspiration of oropharyngeal secretions [2–4]. Despite close anatomic proximity, unique microbial communities colonize nasal and oral cavities owing to differences in nutrient availability, oxygen gradients, epithelial mucosa, and immune microenvironments [5–9].

Acquisition of the upper respiratory tract (URT) microbiome occurs during the perinatal period and continues to develop through adulthood. While there have been numerous comparative and longitudinal studies of gut microbiota alterations across the lifespan [10–12], knowledge regarding URT microbiome development remains nascent. Most studies have focused on adult populations, though microbiome dysbioses in early life have been implicated in shaping long-term health trajectories. Moreover, prior investigations have largely been limited to genus-level characterizations via 16S rRNA gene sequencing without evaluation of microbial functioning [13–16].

One key role of native microbiota is to provide colonization resistance against pathogen colonization. The nasal and oral mucosa are frequently colonized by bacterial, viral, and fungal pathogens, particularly in childhood. Many studies report an association between URT pathogen colonization and various infectious and non-infectious diseases in both children and adults [17–25]. Microbial interference that inhibits pathogen acquisition or overgrowth can occur through a range of microbe-microbial or host-microbial interactions, including via antimicrobial specialized metabolites [17]. Our recent work determined the distribution of microbial biosynthetic gene clusters (BGCs) encoding specialized metabolites across the aerodigestive tract, showing various naso-oral sites harbor distinct bacterial communities and BGCs [18]. Functional studies of interference interactions occurring within upper respiratory microbiomes could provide valuable insight into niche-specific mechanisms of colonization resistance. Of particular relevance is understanding how such interactions influence pathogen carriage and microbiome development in early life when infection risk is high.

To address these gaps in understanding how the trajectory of microbiome maturation and functioning relates to respiratory health, we analyzed microbiome composition and functioning among healthy infant and adult populations. We utilize shotgun metagenomic sequencing of paired nasal and oral samples to assess niche-specific microbiome changes. Across respiratory sites, we examine patterns of microbial community composition, diversity, and species-level taxonomic shifts that vary with age. In addition, we leverage metagenomics to compare the presence of antimicrobial resistance alleles and biosynthetic gene clusters encoding specialized metabolites that may mediate microbial interactions. Lastly, through high-throughput co-culture assays we evaluate pairwise upper respiratory commensal-pathogen interactions. This work informs our knowledge on microbial community functioning at upper respiratory sites that are influenced by aging, with important implications for host health.

## Results

### Nasal and Oral Microbiome Composition and Diversity Vary with Age

We first investigated microbial communities of upper respiratory sites among healthy infants and adults. Non-metric multidimentional scaling (NMDS) of Bray-Curtis and Jaccard dissimilarity values identified differences in the microbial community structure of nasal and oral microbiomes (Figure 1A-D, Table S1). Permutational Multivariate Analysis of Variance (PERMANOVA) demonstrated that, among both age groups, samples clustered by body site with a larger R^2^ value for the adult microbiomes (PERMANOVA, R^2^ = 88%, p.adj<0.001) than for infant microbiomes (PERMANOVA, R^2^ = 79%, p.adj<0.001) (Figure 1A-B, Table S1). Beta diversity dispersion was significantly lower across oral metagenomes relative to nasal metagenomes (Figure 1A-D, Table S1). NMDS ordination showed that centroids were distinct, indicating differences were driven by the effect of body site rather than variation in beta-dispersion [19]. Samples from both upper respiratory sites also clustered by age, with nasal microbiomes demonstrating larger R^2^ values than oral microbiomes (PERMANOVA, R^2^ = 25% and 8%, respectively, p.adj<0.001) (Figure 1C-D, Table S1). Nasal microbiomes differed in beta-dispersion, with NMDS ordination showing that centroids were distinct, indicating differences were driven by the effect of age rather than variation in beta-dispersion. Oral microbiomes did not significantly differ in beta diversity dispersion, indicating differences in microbial community structure at this site were primarily impacted by host age. Analysis of similarities (ANOSIM) complemented these findings (Table S1). Comparisons across alpha diversity metrics identified further differences in upper respiratory microbiomes across age groups (Figure 1E). Among nasal microbiomes, adults demonstrated significantly higher median richness and diversity according to Shannon and Fisher indices. In contrast, infant oral microbiomes had higher richness and diversity, relative to adult microbiomes. Across both age groups, oral microbiomes showed significantly higher median richness and diversity compared to nasal microbiomes (Figure S1).

**Figure 1.**
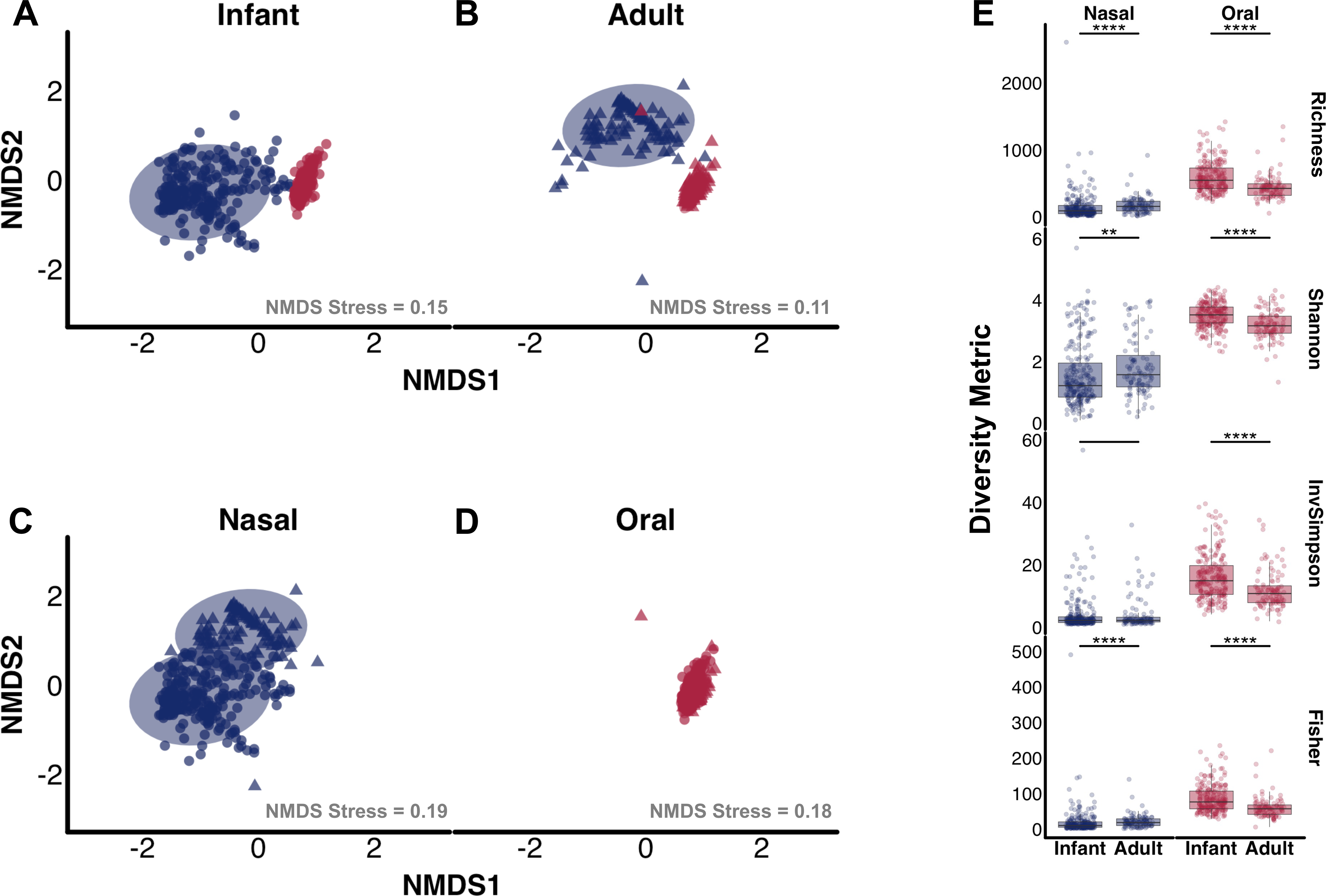
Nasal and Oral Microbial Communities of Infants compared to Adults. Non-metric multidimentional scaling (NMDS) ordination plot of Bray-Curtis dissimilarity indices of (A) infant nasal and oral microbiomes, (B) adult nasal and oral microbiomes, (C) adult and infant nasal microbiomes, and (D) adult and infant oral microbiomes. Shotgun metagenomic sequencing reads were examined from microbiomes normalized to an equal sequencing depth. Points represent individual samples, with circles denoting infant microbiomes and triangles adult microbiomes. Nasal samples are colored blue, oral samples are colored red. Plot stress are listed for each comparison. (E) Boxplots displaying richness, Shannon, Inverse Simpson, and Fisher diversity metrics of nasal (blue) and oral (red) microbiomes from infants compared to adults. Plot is faceted by body site. Boxes show the 25^th^ and 75^th^ percentiles with the median represented by a horizontal line and whiskers showing 1.5 x the interquartile range. Overlayed data points are jittered to avoid overplotting. Significance values * = p < 0.05 and **** = p < 0.001 Mann–Whitney U-test.

Given the observed differences in microbial community structure and diversity, we next compared taxonomic shifts across upper respiratory sites and age groups. Across nasal microbiomes, we identified significant differences in several genera between infants and adults. (Figure 2A, S3). Adult nasal microbiomes had significantly higher relative abundance of *Cutibacterium, Bacteroides, Corynebacterium,* and *Rothia* (median log(abundance) [MLA] = 1.85, 0.19, 0.43, and 0.16, respectively), compared to infant microbiomes (MLA = 0.11, 0.0, 0.03, and 0.06, respectively). Infant nasal microbiomes had significantly higher *Moraxella, Dolosigranulum,* and *Psychrobacter* (MLA = 1.72, 1.04, and 0.003 respectively) compared to adults (MLA = 0.0, 0.003, 0.0, respectively). Accordingly, the majority of infant nasal microbiomes (n=117/229) were dominated (>50% relative abundance) by *Moraxella*, while the majority of adult microbiomes (n=66/100) were dominated by *Cutibacterium*. Species-level differences were reflective of differences among these genera (Figure 2B). For instance, infants had significantly higher relative abundance of four *Moraxella* spp. including the respiratory pathogen *M. catarrhalis* (MLA = 0.65) versus adults (MLA = 0.0). Notably, while most *Streptococcus* spp. were in higher abundance among adults, infants had greater abundance of *S. pneumoniae* (MLA= 0.29) compared to adults (MLA = 0.07). Another common respiratory pathogen, *H. influenzae,* was likewise enriched in infant nasal microbiomes (MLA = 0.05), compared to adults (MLA=0.006).

**Figure 2.**
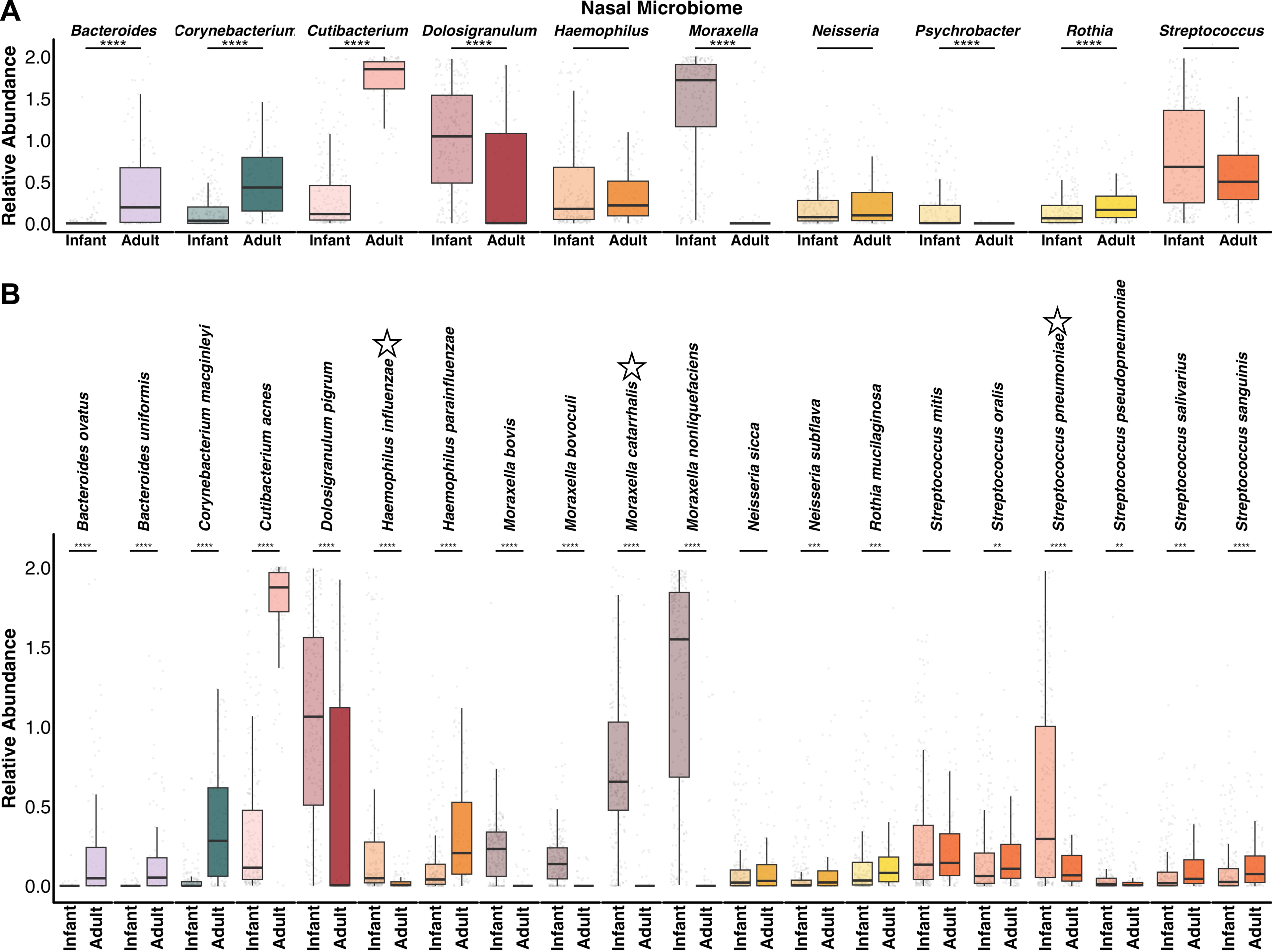
Core Nasal Taxa across Infants and Adults. Boxplots comparing the relative abundance of the ten most abundant (A) nasal genera and (B) top twenty nasal species from these genera between infants and adults. Stars above species names in (B) highlight known respiratory pathogens. Y axes display log + 1 of abundance values. Boxes show the 25^th^ and 75^th^ percentiles with the median represented by a horizontal line and whiskers showing 1.5 x the interquartile range. Overlayed data points (grey) are jittered to avoid overplotting. Significance values * = p < 0.05, ** <0.01, *** = <0.001, and **** = p < 0.0001 Mann–Whitney U-test with FDR correction.

In the oral microbiome, *Streptococcus* was the dominant genus of most infant and adult samples (n=126/212 and n=66/100, respectively) and was not significantly different between age groups (Figure 3A, S2, S3). Of the ten most abundant oral genera, six demonstrated significant differences across age groups (Figure 3A). We found adult oral microbiomes had significantly higher abundance of *Haemophilus* and *Gemella* (MLA= 1.26 and 0.58, respectively), relative to infant microbiomes (MLA= 0.87 and 0.49, respectively). Infants had higher relative abundance of *Neisseria, Rothia, Alloprevotella,* and *Capnocytophaga* (MLA= 1.07, 0.92, 0.33, and 0.28, respectively), compared to adult microbiomes (MLA= 0.46, 0.59, 0.09, and 0.16, respectively). Species-level data revealed, for instance, infants had higher relative abundance of *Rothia aeria* and *Rothia mucilaginosa* compared to adults, but not *Rothia dentocariosa* (Figure 3B). *Streptococcus oralis* and *Streptococcus pneumoniae* were found in similar abundance across infants and adults, while additional *Streptococcus* spp. demonstrated significant differences in relative abundance between age groups. In addition, adults demonstrated significantly higher abundance of *Haemophilus parainfluenzae,* but not *Haemophilus haemolyticus*, relative to infants.

**Figure 3.**
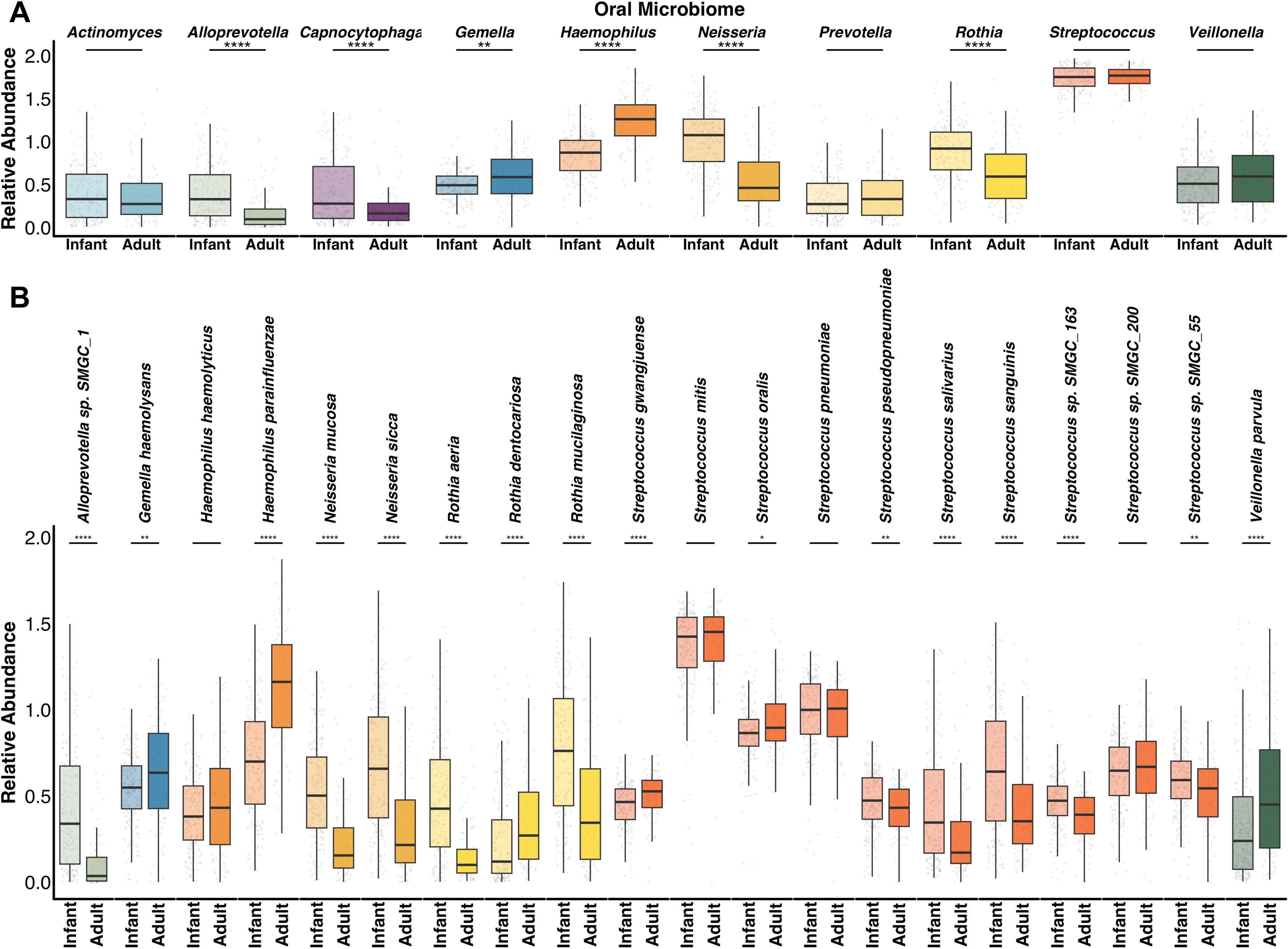
Core Oral Taxa across Infants and Adults. Boxplots comparing the relative abundance of the ten most abundant (A) oral genera and (B) top twenty oral species from these genera between infants and adults. Stars above species names in (B) highlight known respiratory pathogens. Y axes display log + 1 of abundance values. Boxes show the 25th and 75th percentiles with the median represented by a horizontal line and whiskers showing 1.5 x the interquartile range. Overlayed data points (grey) are jittered to avoid overplotting. Significance values * = p < 0.05, ** <0.01, *** = <0.001, and **** = p < 0.0001 Mann–Whitney U-test with FDR correction.

### Antimicrobial Resistance Gene Landscape of Nasal and Oral Microbiomes

Owing to the finding that the nasal and oral cavities of infants and adults harbor distinct microbial communities, we hypothesized that the resistomes of these groups would likewise differ. We aligned metagenomes against the curated Comprehensive Antibiotic Resistance Database (CARD) to determine antimicrobial resistance gene (ARG) presence. We identified reads mapping to protein-coding ARGs from nasal (n=137) and oral (n=309) metagenomes, respectively. ARGs clustered primarily by body site, rather than age group (Figure S4A). Indeed, median ARG abundance from all oral metagenomes was higher than nasal (8.9 vs. 7.6 Fragments Per Kilobase Million [FPKM]), while no significant difference in median abundance across age groups was found (Figure S4B-D). Comparison across body sites identified 153 and 28 ARGs that were differentially abundant in nasal versus oral sites, respectively, while 95 were shared (Figure S4E). The gene ICR-Mc from *M. catarrhalis* was most enriched nasally, while APH(3’)-lla was most unique to the oral cavity (Table S2). While median abundance did not significantly differ with age at either body site, examination of differential ARGs identified unique patterns of enrichment (Figure 4A-B, Table S3). From oral metagenomes, 25 and 33 ARGs were significantly more abundant in infant or adult metagenomes, respectively (Figure 4A). 26 and 65 ARGs were enriched in infant or adult nasal metagenomes, respectively (Figure 4B). Notably, several of the genes from infant nasal metagenomes originated from respiratory pathogens (*H. influenzae, M. catarrhalis*, or *S. pneumoniae*). We therefore evaluated median abundances of ARGs originating from these pathogens. Infant nasal metagenomes contained significantly higher ARGs from *H. influenzae* and *S. pneumoniae*, while adult oral metagenomes had higher *H. influenzae* ARGs (Figure 4C). No differences in *M. catarrhalis* ARG abundance were identified between infants and adults.

**Figure 4.**
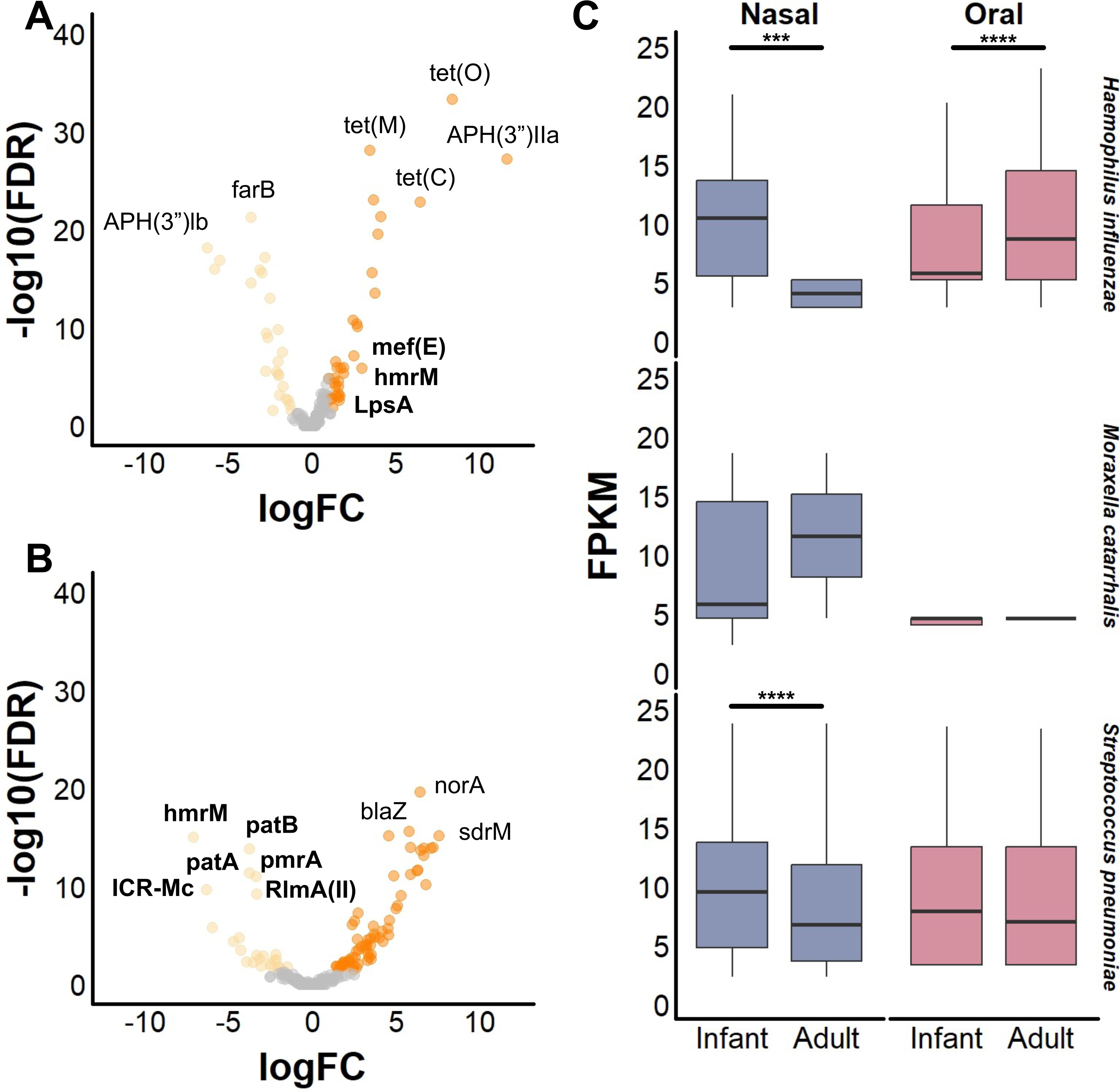
Antimicrobial Resistance Gene Content. Volcano plots comparing antimicrobial resistance genes identified from (A) oral and (B) nasal metagenomes that are differentially abundant between infants (light orange points) and adults (dark orange points), or not significantly different (grey points). Y-axis represents inverse of log of FDR significance values and x-axis represents the log fold-change between groups. FDR cutoff of <0.05 was used. Select genes are labeled, with bold text indicating a gene originated from a respiratory pathogen (H. influenzae, M. catarrhalis, or S. pneumoniae). Metagenomes (n=89 infant nasal, n=212 infant oral, n=48 adult nasal, n=97 adult oral) were subsampled to 500,000 paired reads and aligned to the CARD database. Counts of antimicrobial resistance genes with MAPQ greater or equal to 10 were compared using edgeR. (C) Boxplots of fragments per kilobase per million mapped reads (FPKM) aligning to antimicrobial resistance genes with MAPQ greater or equal to 10 from *H. influenzae*, *M. catarrhalis*, or *S. pneumoniae* between infants and adults. Boxes show the 25^th^ and 75^th^ percentiles with the median represented by a horizontal line and whiskers showing 1.5 x the interquartile range. Significance values * = p < 0.05, ** <0.01, *** = <0.001, and **** = p < 0.0001 Mann–Whitney U-test.

### Distribution of Biosynthetic Gene Clusters Across Adult versus Infant Upper Respiratory Microbiota

We next sought to determine whether the observed shifts in community composition and AMR abundance from infancy to adulthood coincided with changes in BGC abundance. The unique bacterial communities of the upper respiratory tracts harbor distinct BGCs encoding metabolites that are likely to mediate interactions with pathogens and host mucosa [18]. We first examined the abundances of subsampled metagenomic reads aligning to arylpolyene, non-ribosomal polyketide synthetase (NRPS), polyketide synthase (PKS), ribosomally synthesized and post-translationally modified peptide (RiPP), siderophore, terpene, or hybrid BGCs identified from aerodigestive tract bacteria. On average, a greater number of reads from oral microbiomes mapped to BGCs, compared to nasal microbiome reads (p<0.0001), in line with prior findings [18] (Table S4). Infant oral microbiomes possessed the overall highest BGC count, which was significantly higher relative to adult oral microbiomes (p<0.0001, Figure 5A). Infant oral microbiomes had significantly higher arylpolyene, NRPS, PKS, and terpene abundance, compared to adults (Figure 5B, Table S4). Evaluation across BGC classes showed that infant nasal microbiomes had a higher abundance of RiPP and hybrid clusters, while adults had higher arylpolyene, NRPS, PKS, and terpene clusters (Figure 5B, Table S4).

**Figure 5.**
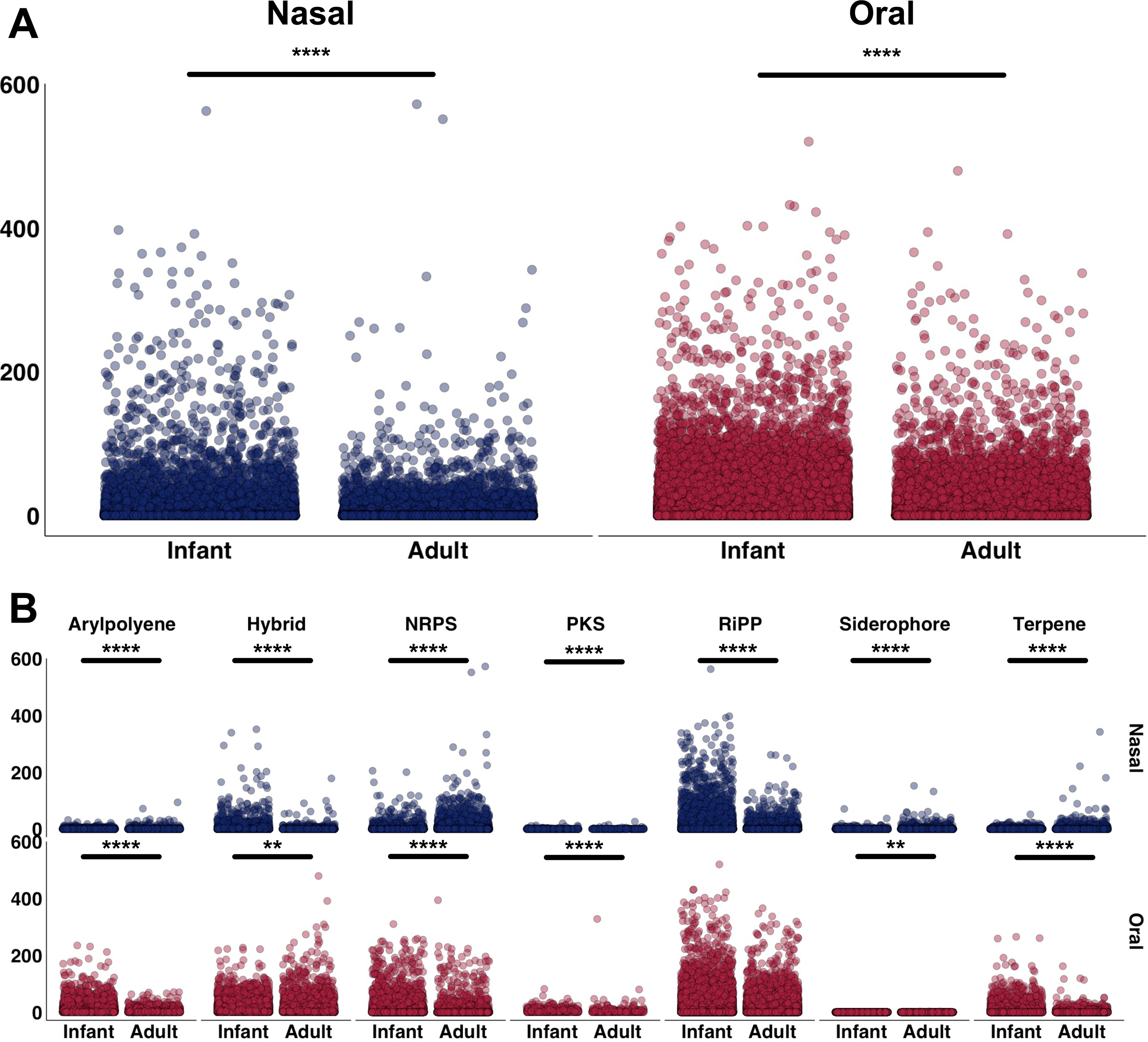
Biosynthetic Gene Cluster Abundance Differs with Age. (A) The plot indicates reads from nasal (blue) and oral (red) infant and adult microbiomes that mapped to BGCs (arylpolyene, NRPS, PKS, RiPP, siderophore, terpene, and hybrid clusters) identified from upper respiratory bacteria (eHOMD habitat listed as “Nasal”, “Nasal,Oral”, or “Oral”). (B) Reads from nasal (blue) and oral (red) infant and adult microbiomes mapping to BGCs from upper respiratory bacteria, grouped by cluster type. All metagenomes were subsampled to 160K reads. 51 infant nasal, 22 adult nasal and 1 adult oral metagenomes were not included in analysis due to low sequencing depth. Normalized reads were pseudoaligned onto an in-house BGC library. NRPS = non-ribosomal peptide synthases, PKS = polyketide synthases, RiPP = ribosomally synthesized and post-translationally modified peptides. Significance value * = p < 0.05, ** = p <0.01, *** = p <0.001, **** = p <0.0001 Yuen Welch’s T-test with 0.001 trim.

We next examined the taxonomic origin of BGCs classes. Infant nasal microbiomes had a higher abundance of RiPPs from *Dolosigranulum* and *Moraxella,* relative to adults (Table S5). Adult nasal microbiomes contained higher average NRPS, PKS, and terpenes from *Corynebacterium*, NRPS from *Cutibacterium*, and NRPS, siderophores, and terpenes from *Staphylococcus*, relative to infants (Table S5). Infant oral microbiomes contained greater average numbers of BGCs from *Neisseria* and *Rothia* compared to adults (Table S5). In particular, arylpolyene, PKS, terpene clusters from *Neisseria* and NRPS from *Rothia* were present in higher abundance, relative to BGCs from these oral taxa in adults (Table S5). Adult oral microbiomes demonstrated higher average *Haemophilus* BGCs belonging to hybrid and RiPP clusters, relative to infants.

### Nasal and Oral Isolates Demonstrate Niche-specific Patterns of Bioactivity

To begin corroborating the role of upper respiratory taxa in mediating colonization resistance, we assessed the capacity of nasal and oral bacteria to inhibit pathogens in a high-throughput co-culture assay [19]. In total, we screened 766 (n=169 individuals) and 301 (n=128 individuals) nasal and oral isolates, respectively, against nine gram-positive, ten gram-negative, and five fungal pathogens. In sum, 25,608 pairwise interactions were evaluated. By examining inhibition across these pathogen types, we found that nasal and oral isolates exhibited unique patterns of inhibition (Figure 6A-B, Table S6-7). We next quantified the average inhibition of strains from each body site versus each pathogen type (i.e. fractional inhibition). We found oral isolates demonstrated higher fractional inhibition of both gram-positive and gram-negative bacterial pathogens (0.49±0.01 and 0.44±0.009, respectively) relative to nasal isolates (0.23±0.005 and 0.11±0.004, respectively), including pathogens of high clinical priority such as *Pseudomonas aeruginosa*, *Acinetobacter baumannii*, and methicillin-resistant *Staphylococcus aureus* (Figure 6C). While gram-negative inhibition by nasal pathogens was overall rare, most isolates did partially or completely inhibit the respiratory pathogen *M. catarrhalis* (Figure 6A). Oral isolates of *Granulicatella* and *Streptococcus* exhibited the highest fractional inhibition of gram-negative and gram-positive pathogens (Figure 6E). Conversely, nasal isolates exhibited higher fractional inhibition of fungal pathogens (0.27±0.007), particularly *Candida* spp. and *Cryptococcus neoformans*, to a greater degree that oral isolates (0.05±0.006) (Figure 6A-C). In particular, nasal *Micrococcus* and *Kocuria* isolates showed high fractional inhibition of fungi (Figure 6E).

**Figure 6.**
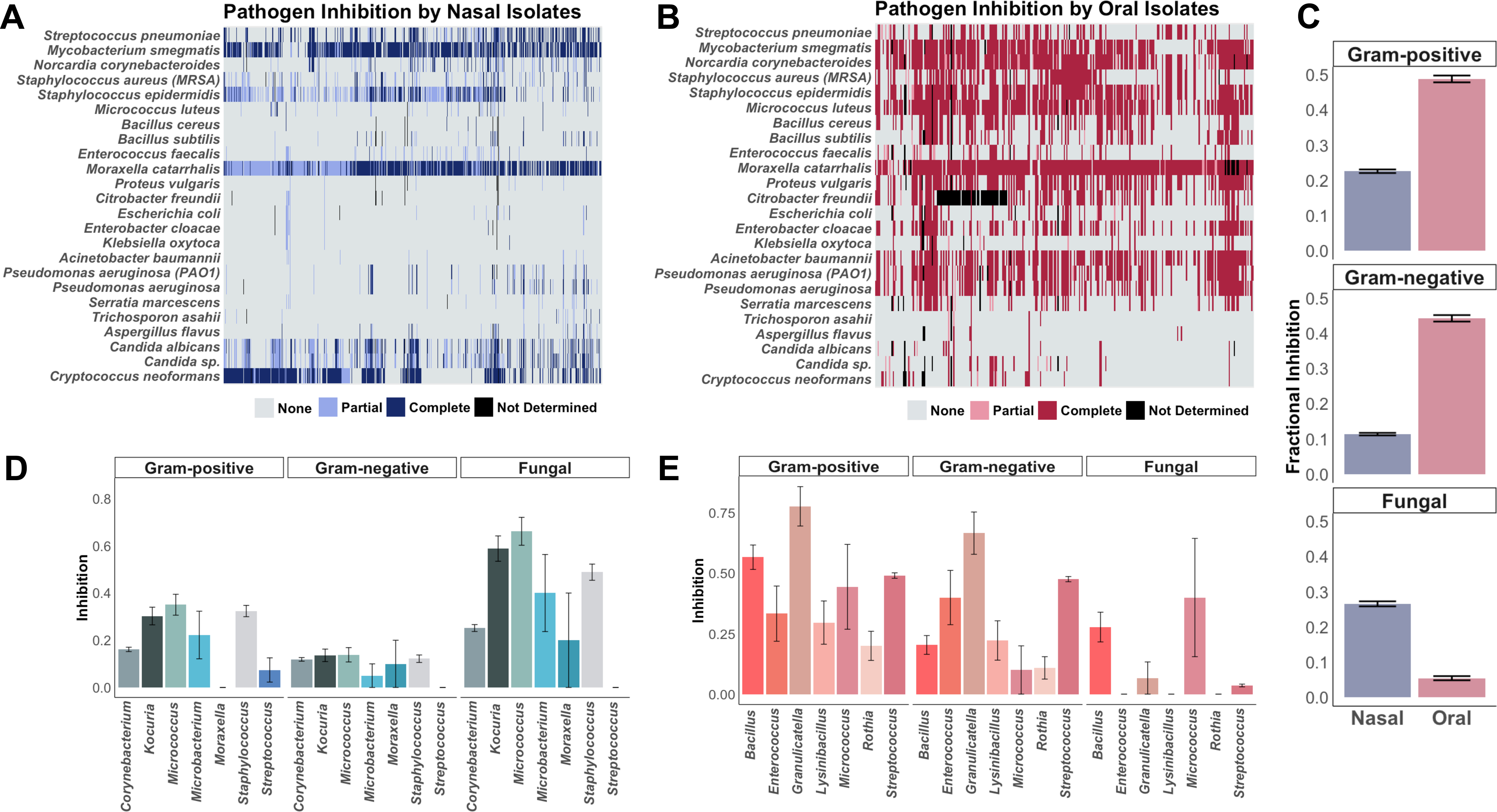
Colonization Resistance Varies across Upper Respiratory Tract Sites. Heatmaps of pathogen inhibition by (A) nasal and (B) oral isolates. Pathogens listed on the y-axis are grouped by gram-positive (top), gram-negative (middle), and fungal (bottom). Inhibition scores (“None”, “Partial”, “Complete”, or “Not Determined”) are colored according to legends inhibition (C) Fractional inhibition scores were calculated for nasal and oral isolates across pathogen types. Fractional inhibition scores for isolates with genus-level taxonomic assignments and are displayed for (D) nasal and (E) oral isolates. Error bars represent standard error of the mean.

## Discussion

While it is becoming increasingly evident that human-associated microbes mediate a multitude of key functions, there is a clear knowledge gap on how respiratory microbiomes contribute to host health across the lifespan. In particular, study of the microbial interactions occurring at mucosal sites has remained limited, but can provide a clearer understanding of the factors shaping microbiome composition. In this work, we utilize shotgun metagenomic sequencing paired with functional pathogen inhibition assays to investigate upper airway microbiome development and its role in pathogen inhibition. To our knowledge, this represents the first large-scale comparison of paired nasal and oral metagenomes from healthy infant and adult populations. Our data identify unique age-related patterns of microbial community composition, diversity, antimicrobial gene content, and colonization resistance across upper respiratory sites.

First, based on shotgun metagenomic sequencing, we find that the microbiota inhabiting nasal and oral cavities are unique among infants and adults, providing further evidence that healthy respiratory microbiota are niche-dependent [13,15]. Comparisons of beta diversity found that the effect of body site was greater than age, in accordance with distinct microbial communities inhabiting each niche. We did identify, however, that nasal microbiota showed greater separation between adults and infants compared to oral flora. This suggests that the nasal and oral microbiomes develop at different rates, with nasal microbiomes having not achieved the same degree of niche differentiation as is present in adults. Prior work that has identified loss of topography between nasal and oral sites from adulthood to elderly age [20]. Together, with our data, this suggests that distinction in upper respiratory microbiota may peak in adulthood. In addition, we show oral microbiomes exhibit high similarity across individuals irrespective of age, whereas nasal microbiomes have greater beta dispersion suggesting more variability across populations. Comparative analyses of nasal-oral microbiomes have shown similar patterns of population-level variation [21,22], possibly owing to differences in inhaled rather than ingested microbes [23], host genetics [24], or influence of environmental changes [25]. Oral communities may exhibit more resilience to exogenous microbes [26], relative to nasal microbiomes, in line with our demonstration of higher fractional inhibition of bacterial pathogens by oral isolates and greater BGC relative abundance.

In terms of diversity, oral microbiomes of infants demonstrate greater richness and taxonomic diversity than those of adults. Previous studies utilizing 16S rRNA gene sequencing of younger individuals (i.e. < 24-months) have reported infants to have lower diversity [27–29]. Here, we report diversity based on metagenomic sequencing that captured non-bacterial microbiome members, providing a more complete view of microbial diversity. In addition, prior comparisons find higher oral microbiome diversity among older children [30], suggesting there is a peak in diversity in childhood that declines in adulthood which warrants further longitudinal study.

Together, our data indicate that early-life oral microbiomes are more diverse than adults, but that these microbial communities have overall low intra-group variation. In contrast, we find nasal microbiomes are less diverse in infancy, providing further evidence that the nasal microbiome is still developing. Previous limited evaluations of nasal microbiota find patterns of diversity differ between children and adults [31,32] with increasing diversity from birth [11,33,34], though factors influencing maturation trajectories remain to be fully defined.

We observed the nasal microbiomes of infants were dominated by two genera - *Moraxella* and *Dolosigranulum* - whereas in most adults, *Cutibacterium* (formerly *Propionibacteria*) had the highest relative abundance. The striking difference in relative abundances of these genera point toward successional patterns of development, wherein infant *Moraxella* (primarily *M. nonliquefaciens*) comes to be replaced by Cutibacterium. Several other genus-level differences were identified, including higher abundance of *Bacteroides*, *Corynebacterium*, and *Rothia* among adults compared to infants. Few large-scale comparisons exist of nasal microbiota across age groups. One study, utilized 16S rRNA gene sequencing to compare microbiomes of healthy children (5-6 years-old [yo]), young adults (18-20 yo) and seniors (50-90 yo) residing in China, finding enrichment of *Moraxella* and *Dolosigranulum* in youth and greater abundance of *Staphylococcus* and *Bacteroides* in adults [35]. We likewise show *Bacteroides* increase with age, but find *Staphylococcus* to be in lower abundance across adult nasal microbiomes. The implications of nasal incursion by *Bacteroides* spp., which are anaerobes, remain understudied but have been linked to chronic rhinosinusitis development [36,37]. The differences in the dominant taxa of adult nasal microbiomes identified by this work compared to our findings may reflect population-level variance due to geography, environmental exposures, and lifestyle factors that merit further investigation. Prior work has also shown increasing abundance of nasal *Corynebacterium* and *Cutibacterium* with age, the latter owing to changes in hormone-dependent sebum production [38]. Notably, we show adult nasal microbiota contain higher relative abundance of several *Streptococcus* spp. compared to infants, including greater incursion of oral species (e.g. *S. oralis*, and *S. salivarius*). Prior work has shown that, relative to adults, elderly individuals likewise have increased abundance of streptococci, suggesting the overlap in nasal and oral microbiomes may begin in adulthood with specific streptococci. Lastly, we also find infant nasal have higher relative abundance of the respiratory pathogens *M. catarrhalis*, *S. pneumoniae*, and *H. influenzae* consistent with culture- and PCR-based studies [39–41]. Oral microbiomes of infants and adults also demonstrated presence of *S. pneumoniae*, suggestive of shared niche-overlap for this pathogen.

In oral microbiomes, we show *Streptococcus* is the most abundant genus across age groups, with *Neisseria* and *Haemophilus* being the next most abundant taxa in infants and adults, respectively. *Neisseria*, in particular *N. sicca* and *N. mucosa*, and *H. parainfluenzae* colonize the shared microenvironment of gingival plaque [42,43], suggesting the latter may outcompete other taxa with age and development of dentition. Prior work has evaluated oral microbiomes of children (8-17 yo) and adults (>18 yo) showing *Streptococcus*, *Haemophilus*, and *Rothia* are the most abundant genera across both groups, along with enrichment of *Neisseria* and *Capnocytophaga* in youth [30], concordant with our results. Additionally, we find infants demonstrated higher relative abundance of *Alloprevotella*, *Capnocytophaga*, and *Rothia*, while adults higher *Gemella* and *Veillonella parvula*. While several streptococci were differentially abundant, we found no differences in the abundance of *S. mitis* across age groups, consistent with this taxon serving as a founding species that colonizes oral cavities from birth [27,44]. Together, the high degree of shared microbiota between infants and adults point toward near-maturity of oral microbiomes by age 24-months.

Unbiased examination of ARG carriage within nasal and oral microbiomes revealed unique patterns across each age groups, concordant with observed differences in community compositions. The oral microbiome has recently become recognized as a resilient reservoir of ARGs, while the resistome of the nasal microbiome remains poorly understood [45–47]. Broadly, we find that body site appeared to have greater influence on ARG abundance, rather than age, with higher median ARG abundance in oral compared to nasal metagenomes. Together with our observation of higher BGC abundance and fractional inhibition of bacterial pathogens by oral isolates, we postulate oral microbiota engage in more frequent interbacterial chemical interference relative to nasal microbiota. Greater competition would also push communities toward evolved resistance to small-molecule inhibitors, potentially explaining why oral microbiomes contained higher ARG abundance [48]. In line with our finding that infant nasal microbiomes contain higher abundance of *S. pneumoniae* and *H. influenzae*, we show higher abundance of ARGs originating from these pathogens in infants compared to adults. In *S. pneumoniae*, this included patA/patB, pmrA, conferring fluoroquinolone resistance, RlmA(II) conferring macrolide and lincosamide resistance, and mef(E) conferring macrolide resistance. *H. influenzae* carried the efflux pump hmrM conferring fluoroquinolone resistance.

We further demonstrate variation in BGC abundance across age groups with distinct patterns at respiratory sites corresponding with taxonomic shifts. Few comparative studies of BGCs from human microbiota have been performed, with only two examining aerodigestive microbiota of adults [18,49]. To our knowledge, this work represents the first comprehensive examination of BGCs from infant URT microbiomes. We show that infant oral microbiomes exhibit the higher relative BGC abundance overall, compared to adult oral microbiomes, with significantly greater abundances of arylpolyene, PKS, and terpenes from *Neisseria* and NRPS from *Rothia*. While the majority of BGC-encoded metabolites remain uncharacterized, *Rothia* spp. have been shown to contain an NRPS that encodes enterobactin, a small-molecule active against *S. aureus* [50]. Our prior work further demonstrates *Rothia* inhibit *M. catarrhalis* through a peptidoglycan endopeptidase [51]. Here, we confirm *Rothia* isolates inhibit a range of gram-negative and gram-positive pathogens, further implicating this commensal in shaping the URT microbiome. Within the nasal microbiome, infants demonstrated greater abundance of RiPPs from *Dolosigranulum* and *Moraxella*, corresponding to enrichment of these genera in infants relative to adults. *Dolosigranulum* bioactivity has been shown to inhibit *S. aureus* and, when paired with *Corynebacterium* spp., inhibit *S. pneumoniae* [52]. Relative to infants, adult nasal microbiomes showed higher abundance of NRPS, PKS, and terpenes from *Corynebacterium*, NRPS from *Cutibacterium*, and NRPS, siderophores, and terpenes from *Staphylococcus*. Prior work has identified these genera to participate in several microbial interactions, including inhibition of *S. aureus* by *Staphylococus lugdunesis* via lugdunin production [53], by *Cutibacterium* acnes via cutimycin [54], and by *S. epidermidis* via epifadin [55]. Among Corynebacteria, siderophore-mediated competition between *Corynebacterium propinquum* and coagulase-negative staphylococci [56], as well as inhibition of *S. pneumoniae* by *C. accolens* and *C. amycolatum* has been demonstrated [57,58]. Accordingly, we here show nasal corynebacterial and staphylococcal isolates inhibit gram-positive pathogens, including *S. pneumoniae*, *S. aureus*, and *S. epidermidis*, as well as *Candida albicans* and *Cryptococcus neoformans*. Taken together, we show that taxonomic differences among infants and adults correspond with altered BGC compositions, which mediate microbial interactions that shape URT microbiomes.

Finally, investigation of the capacity of URT bacteria to inhibit pathogens revealed higher fractional inhibition of gram-negative and gram-positive bacteria by oral commensals, while nasal isolates had higher antifungal activity. Our work marks the first direct comparison of pathogen inhibition by nasal versus oral bacteria. The observation of varied patterns of colonization resistance across body sites lends support to the concept that microbial communities have co-evolved to provide niche-specific defenses. Among nasal isolates, *Micrococcus* and closely-related *Kocuria* demonstrated the highest fractional inhibition of fungal pathogens. Marine sponge-associated isolates from these genera have been shown to produce kocurin, a molecule with anti-staphylococcal but not antifungal activity [59], suggesting influence of host environment on BGC expression or evolution. In contrast, we noted relatively low rates of antifungal activity by oral isolates. Though the oral mycobiome remains underexplored, several studies point toward mutualistic chemicophysical interactions between oral bacteria and *Candida* sp. [60–63]. Thus, oral communities may exhibit a degree of tolerance toward fungi, especially in the setting of restricted fungal virulence that occurs in the oral cavity [64]. Among oral isolates, we find streptococci and *Granulicatella* exhibited high antibacterial activity. Members of the former genus have been noted to inhibit respiratory and cariogenic pathogens through a range of mechanisms, including bacteriocin production, metabolic scavenging, and hydrogen peroxide production [65–67]. The commensal *Granulicatella* belongs to the “nutritionally variant streptococci” and, prior to this work, was unexplored with respect to its antimicrobial potential.

## Conclusions

In summary, we examine upper respiratory microbiome maturation and functioning among healthy infants relative to adults, finding significant differences in the microbial communities inhabiting nasal and oral cavities. The results of this study are limited to comparisons across two age groups, which represent distinct infant and adult populations. Longitudinal sampling of individuals from early life through adulthood is needed to gain further understanding of aerodigestive microbiome development. Moreover, examination of pairwise commensal-pathogen interactions only provides insight into contact-independent mechanisms, with further study needed to examine these patterns in living systems.

This study provides evidence of population-level differences in aerodigestive microbial communities and informs our understanding of microbiome changes occurring with age. Through shotgun metagenomic sequencing, we define the major compositional shifts occurring at each site and identify differences in diversity and species-level relative abundances. We further show antimicrobial resistance genes and biosynthetic gene clusters encoding specialized metabolites vary with age and body site, which likely govern intermicrobial interactions within these microbial communities. Lastly, through functional co-culture assays we evaluate pairwise upper respiratory commensal-pathogen interactions, finding niche-specific patterns of bacterial and fungal pathogen inhibition. Together, our findings advance our understanding of the microbial interactions shaping microbiome composition occurring across respiratory mucosal sites, with important implications for host health from early-life onward.

## Materials and Methods

### Study Design and Sample Collection

The Wisconsin Infant Study Cohort (WISC) is an ongoing observational birth cohort established in 2013 and has been previously described [68]. Participants born between 2013-18 were concurrently recruited and enrolled at two study sites, Marshfield Clinic and LaFarge Medical Clinic (part of Vernon Memorial Healthcare). Collection kits were from a centralized site with rigorous lot tracking. Oral and nasal microbiome specimens used in this study were collected during routine surveillance timepoints between 2015-2021 from enrolled 24-month-old infants. Specimens were collected by trained research coordinators. Oral specimens were collected using a Catch-All Sample Collection Swab to swab the entire area of both the left and right buccal mucosa for 10 seconds per side. After pressing the swab against the cryovial wall for greater than 20 seconds, specimens were stored at -80°C in PowerBead Solution (Qiagen). Nasal specimens were obtained by swabbing (Copan #56750CS01) the anterior nasal passage, immersing the swab in Amies media, and then cryopreserving it until extraction and culture. Written informed consent was obtained from primary guardians. The WISC study was approved by the Human Subjects Committee at the Marshfield Clinic Health System and University of Wisconsin-Madison (IRB approvals: KEI10613 and 2017-1454).

### DNA extraction and Metagenomic Sequencing

DNA extractions of infant nasal and oral samples were carried in a DNAse, UV-treated, and bleach-treated laminar flow hood. Negative DNA extraction controls (n=12 and n=11 for nasal and oral samples, respectively) were processed and sequenced in tandem with study samples. Two mock microbial communities contained either 20 Strain Staggered Mix Genomic Material (ATCC MSA-1003) or 6 Strain Skin Microbiome Genomic Mix (ATCC MSA-1005) were also included as positive controls. Prior to extraction, samples were thawed on ice and 100 μL of storage media was combined with ReadyLyse (Epicenter), Mutanolysin (Sigma), and Lysostaphin (Sigma), followed by incubation at 37°C with shaking for 1 h. Samples were next added to a glass bead tube (Qiagen) and vortexed for 10 min followed by incubation at 65°C with shaking for 30 min. After transferring sample liquid to a sterile tube, MPC protein precipitation reagent (Lucigen) was added. The sample was vortexed and centrifuged for 10 min at 21,300 × *g*. The supernatant was combined with isopropyl alcohol and column purified (Invitrogen PureLink Genomic DNA Extraction Kit) then eluted in 50 μL elution buffer. DNA was quantified using the PicoGreen dsDNA quantification assay kit (Invitrogen). Illumina sequence libraries were prepared using the Nextera XT DNA Library Preparation Kit (Illumina), followed by pooling and sequencing at the University of Minnesota Genomics Center. 2x150 bp paired-end sequencing of pooled samples was performed on the NovaSeq 6000.

### Adult upper respiratory metagenomes

We downloaded the raw paired-end shotgun metagenomic reads from matched buccal mucosa and external naris samples (n=100 participants) collected through the “Healthy Human Subjects” Human Microbiome Project studies (Table S8).

### Metagenomic Sequence Processing

Raw read data from WISC and HMP participants was first preprocessed using fastp v0.21.0 [69] to filter low quality, reads <50 bp, and remove adapters sequences. Filtered reads were processed using Kneaddata v0.8.0 to remove tandem repeats using Tandem Repeat Finder and to filter and remove human DNA sequences by mapping reads to the human genome. Taxonomic identification was assigned using Kraken2 v2.0.8-beta [70] and abundances estimated using Bracken v2.5 [71], with a custom database including complete bacterial, viral, archaeal, fungal, protozoan, and human genomes along with UnivEc ore sequences from RefSeq. The database separated and filtered plasmid sequences, as plasmid sequences are frequently assigned incorrect taxonomies using Kraken2 [72]. Potential contaminant species were identified from negative extraction controls and excluded from further analysis using the prevalence method in decontam v1.10.0 [73]. Reads not annotated at the phylum level, as well as those belonging to phyla present at <1% prevalence were additionally filtered. For WISC participants, we obtained an average of 849,232 nasal and 12,016,295 oral reads of non-human, quality-filtered, paired-end reads per sample. For HMP participants, we obtained an average of 1,360,588 nasal and 9,880,770 oral reads of non-human, quality-filtered, paired-end reads per sample.

### Metagenomic Sequence Analyses and Statistics

Nonparametric multidimensional scaling (NMDS) was performed on Bray-Curtis distances of normalized read counts using the ordinate function of the phyloseq R package (version 1.42.0) [74]. PERMANOVA and ANOSIM were performed on beta diversity distances with 999 permutations using the vegan R package (version 2.6-4) [75]. The betadisper function of the vegan R package (version 2.6-4) was used to compare homogeneity of beta diversity dispersions. Alpha diversity metrics were determined by comparing normalized read counts using the estimate_richness function of the phyloseq R package (version 1.42.0) [74] and compared using the Mann–Whitney U-test with FDR correction. Mean read count pseudoaligned to BGCs was compared using Yuen’s T test with 0.001 trim. Alpha level of significance was set to <0.05 for all statistical tests with p-values of <0.05, <0.01, <0.001, and <0.0001 represented as *, **, ***, and ****, respectively.

### Detection and Analysis of Antimicrobial Resistance Genes

Processed metagenomic sequencing reads were subsampled to a set count (500,000 reads) via seqtk 1.4. Antibiotic resistance genes were identified by alignment of metagenomes against the Comprehensive Antibiotic Resistance Database (CARD) via KMA (v1.4.9) using RGI (v5.1.0) (rgi bwt -read_one -read_two -output_file -clean -local) [76]. Each gene was annotated with Resistance Mechanism and Pathogen of Origin using CARD metadata. The read count abundance was normalized to Fragments Per Kilobase Million (FPKM) by the formula: FPKM = (counts x 1E9) / (gene_length x sum(counts)), where “counts” represents the number of reads mapped, “gene_length” is the length of the corresponding gene. Median FPKM aligned to antimicrobial resistance genes were compared across groups using the using the Mann–Whitney U-test. Differential abundances of resistance genes were calculated using the edgeR package in R (v3.32.1) [77,78].

### Alignment of Metagenomic Reads to Biosynthetic Gene Clusters

Processed metagenomic sequencing reads were subsampled to a set count via seqtk 1.4. Kallisto quant 0.46.0 [79] was used to pseudoalign reads to an index of URT-associated BGC open reading frames (ORFs), as previously described [18]. Briefly, 1,527 bacterial genome sequences were downloaded from the eHOMD V9.03 [80], from which 3,895 BGCs were identified via antiSMASH [81]. Hmmscan from HMMER 3.1b2 [82] was then used to identify ORFs commonly associated with BGCs. Kallisto was used to build a 31 length k-mer index containing all ORFs, except nonbiosynthetic ORFs. All estimated read counts for all ORFs were aggregated into a single estimated read count per BGC.

### Strain Isolation and Identification

Nasal swab storage medium (100 μL) was inoculated onto BHI agar supplemented with 0.1% Tween-80 +/- 0.005% wt/vol lithium mupirocin (Sigma-Aldrich). Oral swab storage medium (100 μL) was inoculated onto BHI agar supplemented with 0.1% Tween- 80 and Mitis-Salivarius agar (BD). All isolation plates were incubated for 5 days at 37°C. Colonies of distinct morphotype (≥2 colonies per plate) were selected from each sample and passaged on the medium from which isolates were obtained at 37°C. Each isolate was inoculated in 3 mL of BHI supplemented with 0.1% Tween-80 and grown overnight at 37°C with shaking. All isolates were stored at −80°C in 20% (vol/vol) glycerol. Bacterial isolates were identified by sequencing the 16S rRNA gene. Briefly, 1 μL of each bacterial isolate liquid culture was used as the template for PCR amplification of the 16S rRNA gene using the universal 27F (5′- AGAGTTTGATCMTGGCTCAG-3′) and 1492R (5′-CGGTTACCTTGTTACGACTT-3′) primers [83].

Amplification of PCR products was verified using electrophoresis with Tris-acetate-EDTA gels (40 mM Tris base, 20 mM acetic acid, 1 mM EDTA disodium salt, 1% [wt/vol] agarose). PCR products were cleaned and underwent Sanger sequencing using the 27F primer at Functional Biosciences (Madison, WI). Isolates were identified to the genus level using the eHOMD 16S rRNA sequence identifier tool [73].

### Co-culture Inhibition Bioassays and Scoring

Pathogen inhibition of nasal and oral strains was assessed using co-culture plate inhibition assays, as previously described [19,84]. Briefly, each isolate was inoculated in 3 mL of BHI supplemented with 0.1% Tween-80 and grown overnight at 37°C with shaking. Liquid cultures were inoculated onto one half of each well in a 24-well plate containing 1.5 mL/well of basic medium agar supplemented with 0.1% Tween-80 (1% soy peptone, 0.5% yeast extract, 0.5% NaCl, 0.1% glucose, 0.1% K2HPO_4_ pH=7.2)[53]. Plates were incubated at 37°C for 7 days. Bacterial and yeast pathogens were grown in 3 mL of BHI overnight at 28°C with shaking. Pathogen cultures were diluted 1:10 and 1 μL of diluted culture was spotted on the center of each well. Spore stocks of filamentous fungi (stored at −80°C) were diluted 1:10 in BHI prior to inoculation. Plates were again incubated at 28°C for 7 days. Pathogen inhibition was scored 0 to 3 as follows: 0 - no inhibition, 1 - partial inhibition, 2 - presence of a zone of inhibition, 3 - total inhibition. Scores for wells with lack of isolate growth, contamination, or overgrowth were not determined. Fractional inhibition was calculated as previously described [84].

### Statistical Analysis and Data Visualization

All statistical analyses were performed in R with specific packages as described in the preceding sections. Graphics were generated using ggplot2[85] and BioRender.com.

## Acknowledgments

The authors thank the University of Minnesota Genomics Center for metagenomic sequencing services and technical support. We thank Reed Stubbendieck (Oklahoma State University) for assistance in biosynthetic gene cluster analyses and helpful discussions. We thank Timothy Murphy (The State University of New York) for the gift of *M. catarrhalis* strain O35E. Special thanks to WISC study team members for overseeing participant enrollment, visits and sample collections: Casper Bendixsen, PhD (Marshfield Clinic site PI), Kathrine Barnes, MS, MPH, CPH (Marshfield Clinic site Project Manager), Tara Johnson, RN (Marshfield Clinic lead study coordinator). We thank James DeLine, MD (LaFarge Medical Clinic site PI), Gretchen Spicer, CPM, LM, Sheri Hammond, BBA (LaFarge Medical Clinic study coordinators), and Brent Olsen (Marshfield Clinic) for oversight and management of participant data. We are grateful to the study families for their time and commitment.

This work was supported by grants from the National Institutes of Health (U19AI142720 [C.R.C, L.R.K], NIAID F30AI169759 [S.Z], NCATS TL1TR002375 [S.Z], NIAID T32AI055397 [M.H.S], NIAMS F31AR079846 [M.H.S], NIAID U19AI104317 [J.E.G, C.M.S], UL1TR000427 [J.E.G, C.M.S]), the University of Wisconsin School of Medicine and Public Health Wisconsin Partnership Program (C.M.S, J.E.G), and the Jarislowsky Foundation (C.R.C.). The funders had no role in study design, data collection and interpretation, or the decision to submit the work for publication.

## Author contributions

SZ, CMS, JEG, LRK, CRC conceived and designed the analyses. SZ wrote the original manuscript draft. SZ, MHS, SS, TCD performed the experiments, SZ and MHS analyzed data, and SZ, MHS, SS, TCD, CMS, JEG, LRK, and CRC edited the manuscript draft.

## Declaration of interests

We declare that there are no conflicts of interest.

## Data availability

Raw metagenomic sequences generated for this work are deposited in the Short Read Archive under BioProject accession PRJNA1078287. Adult raw metagenome reads were downloaded from the iHMP data portal (https://portal.hmpdacc.org/). Genome sequences used for BGC prediction were downloaded from the eHOMD (http://www.ehomd.org/) and scripts to replicate BGC analyses are available here: https://github.com/reedstubbendieck/adt_bgcs. All other scripts, raw data, and derived data necessary to replicate this work are available here: https://github.com/szelasko1/InfantAdultRespiratoryMicrobiome_paper.

## Supplementary Figure Legends

**Figure S1. Alpha Diversity of Nasal and Oral Microbiomes by Age Group**. Boxplots displaying richness, Shannon, Inverse Simpson, and Fisher diversity metrics of nasal (blue) and oral (red) microbiomes from infants compared to adults. Plot is faceted by age group and comparisons are across body site. Boxes show the 25^th^ and 75^th^ percentiles with the median represented by a horizontal line and whiskers showing 1.5 x the interquartile range. Overlayed data points are jittered to avoid overplotting. Significance values * = p < 0.05 and **** = p < 0.001 Mann–Whitney U-test.

**Figure S2. Distribution of Taxa in Infant and Adult Upper Respiratory Tract Communities.** (A) Variation in bacterial relative abundances of infant nasal (top, n=229) and oral (bottom, n=212) microbiomes at the genus level. (B) Variation in bacterial relative abundances of adult nasal (top, n=100) and oral (bottom, n=100) microbiomes at the genus level. Samples ordered by abundance of top genus per body site and age group. Stacked bars arranged according to genus-level abundance and colored per legends.

**Figure S3. Upper Respiratory Taxa across Infants and Adults.** Boxplots displaying the log of relative abundance of the top ten most abundant nasal versus oral genera across (A) infants and (A) adults. Boxes show the 25^th^ and 75^th^ percentiles with the median represented by a horizontal line and whiskers showing 1.5 x the interquartile range. Overlayed data points (grey) are jittered to avoid overplotting. Significance values * = p < 0.05, ** <0.01, *** = <0.001, and **** = p < 0.0001 Mann–Whitney U-test with FDR correction.

**Figure S4. Antimicrobial resistance genes across metagenomes.** (A) Heatmap displays z-score values of fragments per kilobase per million mapped reads (FPKM) aligning to antimicrobial resistance genes with MAPQ greater or equal to 10 (rows) across nasal and oral metagenomes from infants and adults (columns). Both rows and columns are hierarchically clustered. (B) Boxplots of FPKM aligning to antimicrobial resistance genes with MAPQ greater or equal to 10 between body sites, (C) between age groups, and (D) between age groups faceted by body site. Boxes show the 25^th^ and 75^th^ percentiles with the median represented by a horizontal line and whiskers showing 1.5 x the interquartile range. Significance values **** = p < 0.0001 Mann– Whitney U-test. (E) Volcano plots comparing antimicrobial resistance genes identified to be differentially abundant between all nasal (blue points) and oral (red points) metagenomes, or not significantly different (grey points). Y-axis represents inverse of log of FDR significance values and x-axis represents the log fold-change between groups. FDR cutoff of <0.05 was used. Metagenomes (n=89 infant nasal, n=212 infant oral, n=48 adult nasal, n=97 adult oral) were subsampled to 500,000 paired reads. Normalized reads were aligned to the CARD database. Cells are colored according to abundance of FPKM, with higher values colored in grey and lower values in white. Columns are annotated with respect to body site and age group, colored per legends. Rows are annotated with respect to mechanism of resistance, colored per legends.

**Table S1. Beta diversity and dispersion metrics.** (Top) ANOSIM R statistic and p-value with Benjamini-Hochberg correction and (bottom) PERMANOVA R^2^ effect size, p-value with Benjamini-Hochberg correction, and p-value from beta dispersion comparisons of Bray-Curtis dissimilarity values of nasal and oral microbial communities of adults and infants. Permutations = 999. Shotgun metagenomic sequencing reads were examined from microbiomes normalized to an equal sequencing depth. NS = not significant, * = <0.05, ** = <0.01, *** = <0.001

**Table S2. Differentially abundant antimicrobial resistance genes by body site.** Genes identified to be differentially abundant between nasal and oral microbiomes used to create Figure S4E.

**Table S3. Differentially abundant antimicrobial resistance genes by age and body site.** Genes identified to be differentially abundant between infants and adults used to create Figure 3 A-B. Body site is listed in far right column.

**Table S4. Mean BGC Counts.** Top: Average counts of arylpolyene, NRPS, PKS, RiPP, siderophore, terpene, hybrid, other, and total BGCs across nasal (left) and oral (right) metagenomes from infants and adults. Middle: Comparison of average counts of all BGCs between (left) infant nasal and oral metagenomes and adult nasal and oral metagenomes (right). Bottom: Comparison of total BGCs from key taxa across nasal (left) and oral (right) metagenomes from infants and adults. Significance value * = p-value < 0.05, ** = p-value <0.01, *** = p-value <0.001, Yuen Welch’s T-test.

**Table S5. Mean BGC Counts by Taxa and cluster type.** Average counts of BGCs from key taxa by cluster type between age groups. Significance value * = p-value < 0.05, ** = p-value <0.01, *** = p-value <0.001, Yuen Welch’s T-test.

**Table S6. Results of Co-culture Bioactivity Assays with Nasal Isolates.**

**Table S7. Results of Co-culture Bioactivity Assays with Oral Isolates.**

**Table S8. HMP Adult Microbiome Sample Manifest.**

## Notes

https://github.com/szelasko1/InfantAdultRespiratoryMicrobiome_paper

### Competing Interest Statement

The authors have declared no competing interest.

