## Supplemental Figures S1-4, Table S1, S4, S5 for "Upper respiratory microbial communities of healthy populations are shaped by niche and age"

**Figure S1**

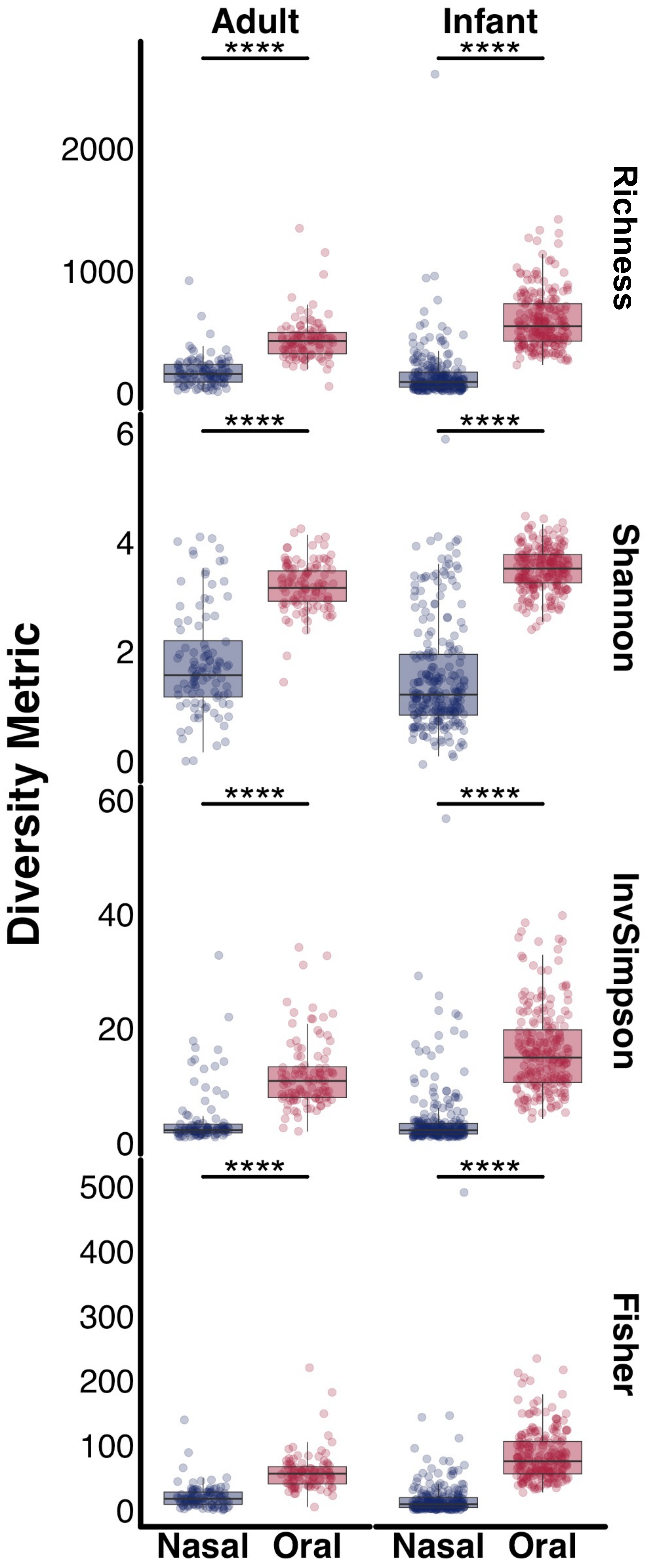

Adult
Infant
****
****
****
****
****
****
****
****
Richness
Shannon
InvSimpson
Fisher
Diversity Metric
2000
1000
0
6
4
2
0
60
40
20
0
500
400
300
200
100
0
Nasal
Oral
Nasal
Oral

### Figure S2

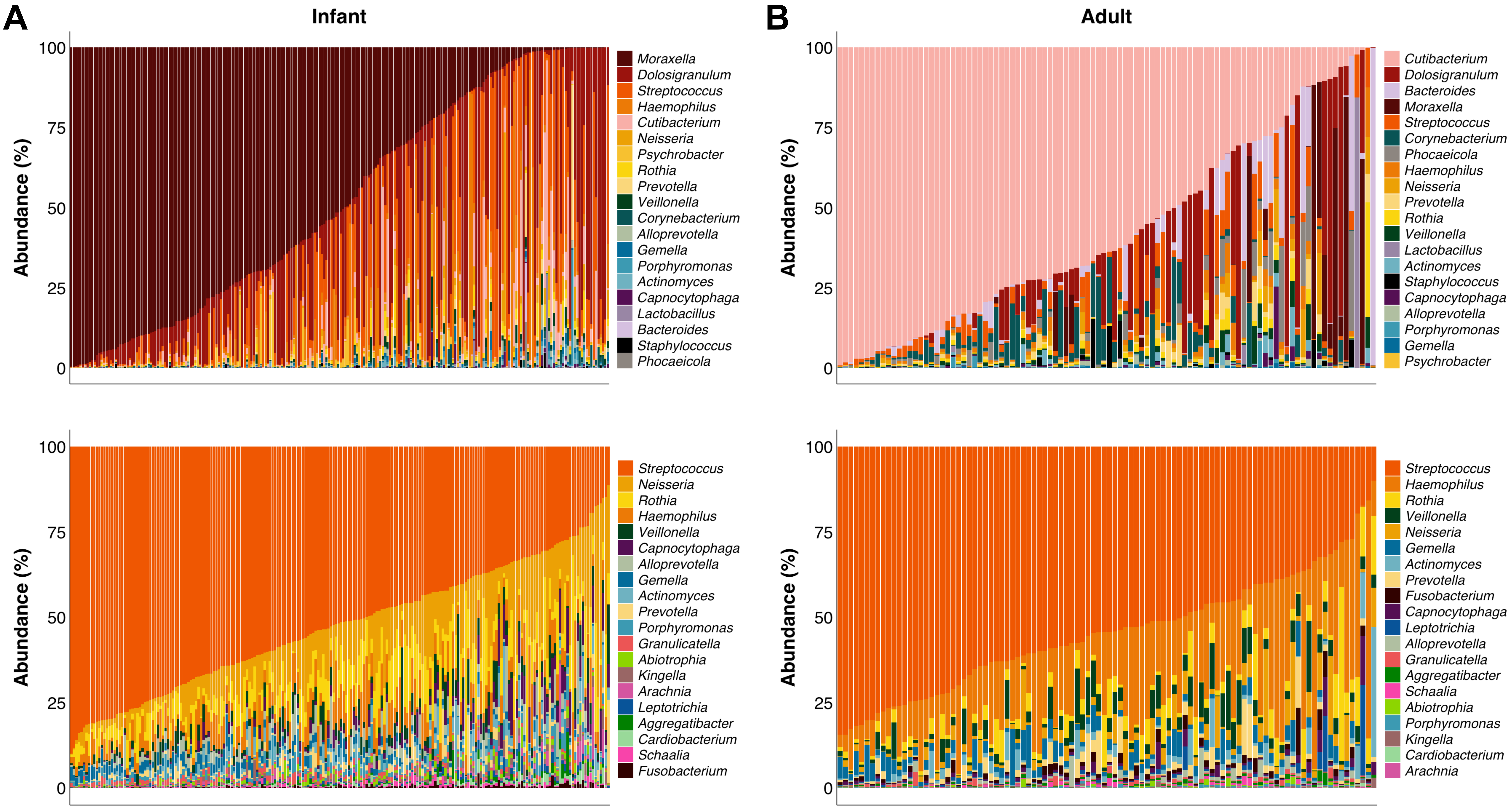
A
Infant
B
Adult
Abundance (%)
0
25
50
75
100
Moraxella
Dolosigranulum
Streptococcus
Haemophilus
Cutibacterium
Neisseria
Psychrobacter
Rothia
Prevotella
Veillonella
Corynebacterium
Alloprevotella
Gemella
Porphyromonas
Actinomyces
Capnocytophaga
Lactobacillus
Bacteroides
Staphylococcus
Phocaeicola
Cutibacterium
Dolosigranulum
Bacteroides
Moraxella
Streptococcus
Corynebacterium
Phocaeicola
Haemophilus
Neisseria
Prevotella
Rothia
Veillonella
Lactobacillus
Actinomyces
Staphylococcus
Capnocytophaga
Alloprevotella
Porphyromonas
Gemella
Psychrobacter
Streptococcus
Neisseria
Rothia
Haemophilus
Veillonella
Capnocytophaga
Alloprevotella
Gemella
Actinomyces
Prevotella
Porphyromonas
Granulicatella
Abiotrophia
Kingella
Arachnia
Leptotrichia
Aggregatibacter
Cardiobacterium
Schaalia
Fusobacterium
Streptococcus
Haemophilus
Rothia
Veillonella
Neisseria
Gemella
Actinomyces
Prevotella
Fusobacterium
Capnocytophaga
Leptotrichia
Alloprevotella
Granulicatella
Aggregatibacter
Schaalia
Abiotrophia
Porphyromonas
Kingella
Cardiobacterium
Arachnia

### Figure S3

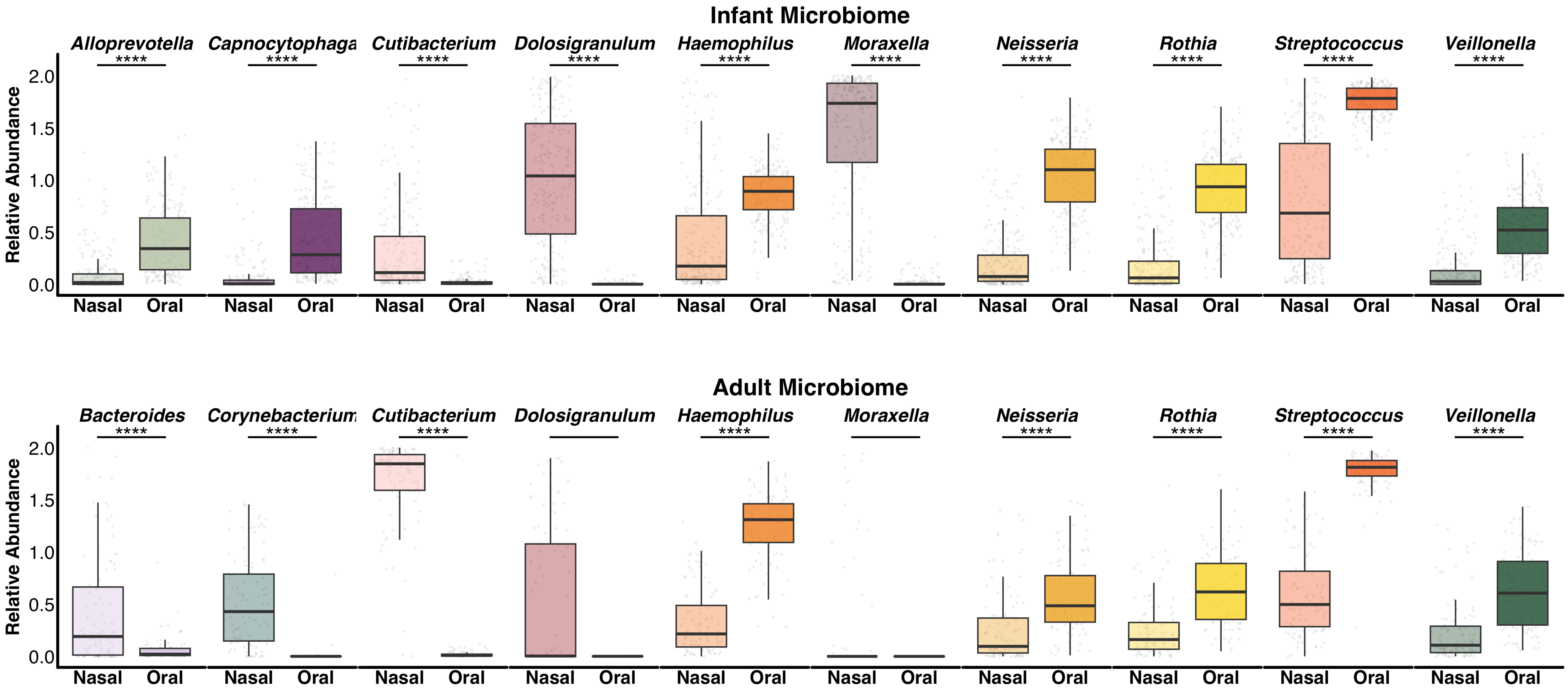

Infant Microbiome
Alloprevotella
Capnocytophaga
Cutibacterium
Dolosigranulum
Haemophilus
Moraxella
Neisseria
Rothia
Streptococcus
Veillonella
****
Relative Abundance
2.0
1.5
1.0
0.5
0.0
Nasal
Oral
Adult Microbiome
Bacteroides
Corynebacterium
Cutibacterium
Dolosigranulum
Haemophilus
Moraxella
Neisseria
Rothia
Streptococcus
Veillonella

Figure S4

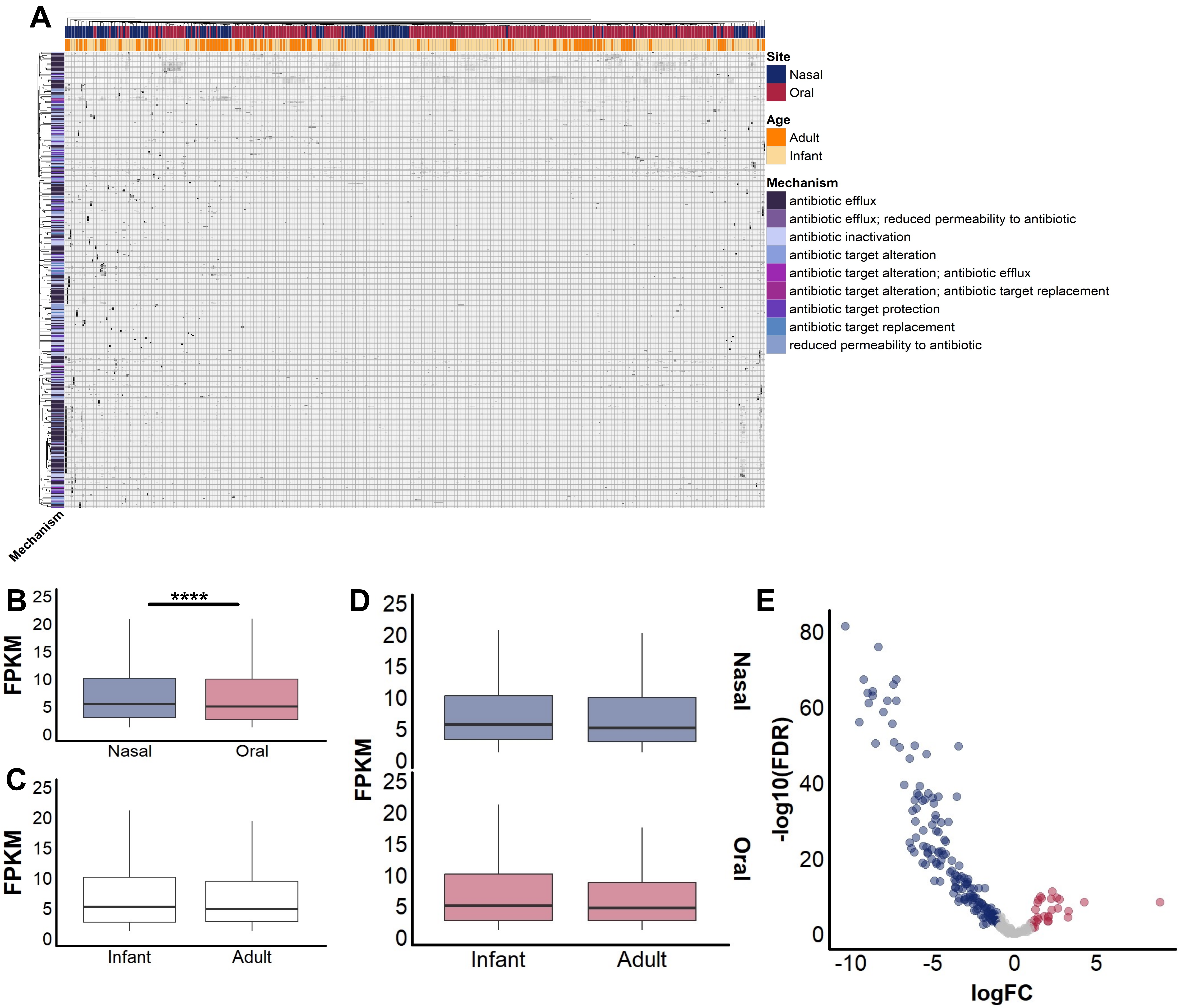
A
Site
Nasal
Oral
Age
Adult
Infant
Mechanism
antibiotic efflux
antibiotic efflux; reduced permeability to antibiotic
antibiotic inactivation
antibiotic target alteration
antibiotic target alteration; antibiotic efflux
antibiotic target alteration; antibiotic target replacement
antibiotic target protection
antibiotic target replacement
reduced permeability to antibiotic
Mechanism
B
****
FPKM
25
20
15
10
5
0
Nasal
Oral
C
FPKM
25
20
15
10
5
0
Infant
Adult
D
FPKM
25
20
15
10
5
0
Nasal
Oral
Infant
Adult
E
-log10(FDR)
80
60
40
20
0
-10
-5
0
5
logFC

#### Table S1

| ANOSIM | | | | | |
|---|---|---|---|---|---|
| | | **Bray-Curtis** | | **Jaccard** | |
| | | R | P.adj | R | P.adj |
| Adult Oral | Adult Nasal | 0.8779 | *** | 0.6488 | *** |
| Infant Oral | Infant Nasal | 0.7885 | *** | 0.7259 | *** |
| Adult Nasal | Infant Nasal | 0.6306 | *** | 0.3449 | *** |
| Adult Oral | Infant Oral | 0.1845 | *** | 0.1748 | *** |

Permutations = 999.
R = ANOSIM R, P.adj = P-value with BH correction
NS = not significant, * = <0.05, ** = <0.01, *** = <0.001

| PERMANOVA | | | | | | | |
|---|---|---|---|---|---|---|---|
| | | **Bray-Curtis** | | | **Jaccard** | | |
| | | $R^2$ | P.adj | B | $R^2$ | P.adj | B |
| Adult Oral | Adult Nasal | 0.48 | *** | * | 0.26 | *** | *** |
| Infant Oral | Infant Nasal | 0.38 | *** | *** | 0.28 | *** | *** |
| Adult Nasal | Infant Nasal | 0.25 | *** | *** | 0.08 | *** | NS |
| Adult Oral | Infant Oral | 0.08 | *** | NS | 0.07 | *** | ** |

Permutations = 999.
P = PERMANOVA P-value with BH correction, $R^2$ = PERMANOVA effect size, B = betadisper P-value
NS = not significant, * = <0.05, ** = <0.01, *** = <0.001

### Table S4

| Mean reads mapping to ADT BGCs from metagenomes | | | | | | |
|---|---|---|---|---|---|---|
| | **Infant Nasal** | **Adult Nasal** | **P value** | **Infant Oral** | **Adult Oral** | **P value** |
| Arylpolyene | 0.1013621 | 0.1581350 | **** | 2.3991372 | 0.9408844 | **** |
| NRPS | 0.8200176 | 3.5093689 | **** | 2.406664 | 1.785770 | **** |
| PKS | 0.7459728 | 2.2812048 | **** | 1.0586320 | 0.6384799 | **** |
| RiPP | 0.5414206 | 0.2459828 | **** | 2.245408 | 2.261341 | NS |
| Siderophore | 0.01914541 | 0.31370781 | **** | 0.0144010 | 0.0063126 | ** |
| Terpene | 0.1973209 | 0.7517247 | **** | 1.5175472 | 0.4548382 | **** |
| Hybrid | 1.4805241 | 0.5113989 | **** | 11.78865 | 10.87360 | ** |
| Other | 0.2333258 | 2.0677877 | **** | 2.773754 | 2.529911 | NS |
| **Total** | **0.4644351** | **0.6427419** | **** | **2.321541** | **1.914613** | **** |

| Mean reads mapping to ADT BGCs from metagenomes | | | | | | |
|---|---|---|---|---|---|---|
| | **Infant Nasal** | **Infant Oral** | **P value** | **Adult Nasal** | **Adult Oral** | **P value** |
| **Total** | **0.4644351** | **2.321541** | **** | **0.6427419** | **1.914613** | **** |

### Table S5

| Mean reads mapping to ADT BGCs from metagenomes | | | | | | |
|---|---|---|---|---|---|---|
| | **Infant Nasal** | **Adult Nasal** | **P value** | **Infant Oral** | **Adult Oral** | **P value** |
| *Corynebacterium* | 3.693553 | 9.353031 | **** | 0.963824 | 0.546378 | **** |
| *Cutibacterium* | 0.4638403 | 10.5399613 | **** | 0.02551800 | 0.08359467 | ** |
| *Dolosigranulum* | 52.77536 | 17.94500 | **** | 0.115566 | 0.085000 | NS |
| *Haemophilus* | 1.0389980 | 0.1895384 | **** | 2.213906 | 6.573915 | **** |
| *Moraxella* | 49.37198 | 3.66500 | **** | 0.01650943 | 0.0000000 | * |
| *Neisseria* | 0.1756493 | 0.2565513 | **** | 5.372873 | 1.761986 | **** |
| *Rothia* | 0.3918197 | 0.7773993 | ** | 8.299404 | 6.067257 | **** |
| *Staphylococcus* | 0.04543505 | 0.96896470 | **** | 0.000448761 | 0.00163265 | *** |
| *Streptococcus* | 0.6935880 | 0.1055103 | **** | 4.305571 | 4.300334 | NS |

| Mean reads mapping to *Dolosigranulum* BGCs | | | |
|---|---|---|---|
| | **Infant Nasal** | **Adult Nasal** | **P value** |
| RiPP | 52.77536 | 17.94500 | **** |

| Mean reads mapping to *Moraxella* BGCs | | | |
|---|---|---|---|
| | **Infant Nasal** | **Adult Nasal** | **P value** |
| RiPP | 49.37198 | 3.66500 | **** |

| Mean reads mapping to *Corynebacterium* BGCs | | | |
|---|---|---|---|
| | **Infant Nasal** | **Adult Nasal** | **P value** |
| NRPS | 6.250881 | 11.976594 | **** |
| PKS | 4.759936 | 14.411120 | **** |
| RiPP | 5.194060 | 4.226639 | NS |
| Terpene | 1.310834 | 4.724415 | **** |

| Mean reads mapping to *Cutibacterium* BGCs | | | |
|---|---|---|---|
| | **Infant Nasal** | **Adult Nasal** | **P value** |
| NRPS | 0.7217897 | 16.2726419 | **** |
| RiPP | 0.3033237 | 6.8014020 | **** |

| Mean reads mapping to *Staphylococcus* BGCs | | | |
|---|---|---|---|
| | **Infant Nasal** | **Adult Nasal** | **P value** |
| NRPS | 0.1096242 | 2.3500505 | **** |
| RiPP | 0.06633063 | 1.70418282 | **** |
| Siderophore | 0.01874662 | 0.31247128 | **** |
| Terpene | 0.04552363 | 0.58346032 | **** |

| Mean reads mapping to *Neisseria* BGCs | | | |
|---|---|---|---|
| | **Infant Oral** | **Adult Oral** | **P value** |
| Arylpolyene | 8.822316 | 2.926161 | **** |
| PKS | 5.192989 | 2.179814 | **** |
| Terpene | 3.0774093 | 0.9775381 | **** |

| Mean reads mapping to *Rothia* BGCs | | | |
|---|---|---|---|
| | **Infant Oral** | **Adult Oral** | **P value** |
| Arylpolyene | NA | NA | NA |
| NRPS | 8.152444 | 6.027143 | *** |
| PKS | 6.985849 | 12.540000 | NS |

| Mean reads mapping to *Haemophilus* BGCs | | | |
|---|---|---|---|
| | **Infant Oral** | **Adult Oral** | **P value** |
| Hybrid | 19.90894 | 58.60600 | **** |
| RiPP | 0.4072894 | 1.3246152 | **** |
